# Influence of the TorD signal peptide chaperone on Tat-dependent protein translocation

**DOI:** 10.1101/2021.03.19.436180

**Authors:** Umesh K. Bageshwar, Antara DattaGupta, Siegfried M. Musser

**Author notes:** Corresponding author: Siegfried Musser.

## Abstract

The twin-arginine translocation (Tat) pathway transports folded proteins across energetic membranes. Numerous Tat substrates contain co-factors that are inserted before transport with the assistance of redox enzyme maturation proteins (REMPs), which bind to the signal peptide of precursor proteins. How signal peptides are transferred from a REMP to a binding site on the Tat receptor complex remains unknown. Since the signal peptide mediates both interactions, possibilities include: i) a coordinated hand-off mechanism; or ii) a diffusional search after REMP dissociation. We investigated the binding interaction between substrates containing the TorA signal peptide (spTorA) and its cognate REMP, TorD, and the effect of TorD on the *in vitro* transport of such substrates. We found that *Escherichia coli* TorD is predominantly a monomer at low micromolar concentrations (dimerization *K*_*D*_> 50 µM), and this monomer binds reversibly to spTorA (*K*_*D*_≈ 1 µM). While TorD binds to membranes (*K*_*D*_≈ 100 nM), it has no apparent affinity for Tat translocons and it inhibits binding of a precursor substrate to the membrane. TorD has a minimal effect on substrate transport by the Tat system, being mildly inhibitory at high concentrations. These data are consistent with a model in which the REMP-bound signal peptide is shielded from recognition by the Tat translocon, and spontaneous dissociation of the REMP allows the substrate to engage the Tat machinery. Thus, the REMP does not assist with targeting to the Tat translocon, but rather temporarily shields the signal peptide.

## INTRODUCTION

The Tat machinery is mechanistically unique in that it transports folded proteins across energetic membranes without collapsing ion gradients. It is the only known protein transport system for which a proton motive force (pmf) is essential for all substrates transported (1-3). In prokaryotes, the Tat machinery transports proteins across the cytoplasmic membrane from the cytoplasm to the periplasm (4-8). Many bacterial Tat substrates are co-factor containing redox proteins. These co-factors, such as molybdopterins or metal centers, are integrated into proteins during folding in the cytoplasm prior to transport (9). Though it remains uncertain exactly how co-factor insertion is ensured before transport, it is clear that redox enzyme maturation proteins (REMPs) play a critical role (10-13). These REMPs generally exhibit selective and specific binding interactions with a single cognate signal peptide, and act as dedicated chaperones (14-16). In addition to their role in redox cofactor assembly and insertion, activities ascribed to REMPs include assistance with protein folding, subunit assembly, proofreading, proteolytic protection, and membrane/translocase targeting (10,12,13,17-21).

In *Escherichia coli*, a functional Tat system minimally consists of three membrane proteins, TatA (or TatE), TatB and TatC (8,22-25). A receptor complex, consisting of an oligomer of TatB and TatC, and likely TatA, binds the signal peptide of transport cargos (26-33). In the presence of a pmf, the mature domain of the protein is conveyed across the membrane with the assistance of additional recruited TatA molecules (34,35). TatC, the largest of the three proteins, contains a large open groove (36,37) that accommodates the signal peptide in a hairpin configuration that extends about halfway across the membrane (33). How the signal peptide transitions from the cytoplasm into this groove, and whether this groove is directly exposed to the membrane interior or directly accessible from the cytoplasmic milieu remain open questions. One model is that the precursor protein binds to the membrane surface via its signal peptide and then diffuses laterally to the Tat receptor complex (38). However, this model is challenged by multiple studies that place the signal peptide binding site on the inside of the receptor oligomer (32,39-41).

At least 8 proteins exported from the *E. coli* cytoplasm by the Tat machinery have signal peptides that bind to a REMP (17,42). While signal peptide binding affinities for REMPs are in the low micromolar to high nanomolar range (43-46), these are potentially significantly modulated by high affinity REMP interactions with the folded mature domain of the cognate protein (47,48). How the signal peptide is able to efficiently locate the signal peptide binding site on the Tat translocon in the presence of REMPs with similar affinities remains unresolved, though a GTP binding site suggests that affinities could be modulated by nucleotide hydrolysis (20,45,49). One hypothesis is that REMPs could deliver their cognate transport cargos to the Tat receptor complex via a coordinated hand-off type mechanism (50). This scenario predicts binding interactions between REMPs and the Tat machinery, consistent with TatB and TatC interactions for DmsD, the REMP for DmsA (51,52). Alternatively, the REMP could be released from the signal peptide within the cytoplasmic milieu, and the free signal peptide could find the Tat translocon in the same manner used by REMP-independent substrates, i.e., by a diffusional search. The absence of a role for DmsD in DmsA translocation *in vivo* supports this model (53).

The oligomerization state of REMPs can potentially influence their various biochemical activities. The X-ray structure of TorD, the REMP for trimethylamine N-oxide reductase (TorA), reveals an extreme domain-swapped dimer (54,55). However, monomeric structures were observed for other TorD family chaperones (14,56-58). Whether the monomer, dimer, or both forms are involved in the various activities ascribed to REMPs remains unresolved. Only the dimer of TorD exhibits GTPase activity, albeit with very low catalytic specificity (20), suggesting that only this oligomeric form could be used for targeting TorA to the Tat translocon followed by GTP-dependent release. Monomeric TorD is sufficient to bind the TorA signal peptide, and both full- and mature-length TorA (45,47), indicating strong interactions of monomeric TorD with both the signal peptide and the mature domain of TorA.

The involvement of REMPs in translocon targeting remains poorly addressed. Here, we examined the *in vitro* binding interactions of TorD with fluorescent proteins fused to the TorA signal peptide (spTorA), and then examined the binding interactions and transport of a REMP/precursor complex with Tat-containing inverted membrane vesicles (IMVs). Monomeric TorD is sufficient for strong signal peptide interactions, yet it does not bind to Tat translocons, or enhance translocon binding of a protein fused to spTorA.

## RESULTS

### *E. coli* TorD is predominantly a monomer

The objective of this study was to examine the influence of TorD on the *in vitro* Tat transport of a folded protein fused to spTorA. Proteins (Table 1 and Figure 1) were overproduced in *E. coli*, and purified by Ni-NTA and size-exclusion chromatographies (Experimental Procedures). We first determined the oligomerization state of *E. coli* TorD (purified and used herein as the 6xHis tagged version TorD-H6). Native gel electrophoresis of TorD (Figure 2A) revealed that the Ni-NTA purified protein predominantly exists in the monomeric form, and trace amounts of higher-order oligomers were eliminated by the addition of the reducing agent β-mercaptoethanol (βME), indicating disulfide-linked oligomers. At the higher concentrations used during size-exclusion chromatography of the Ni-NTA purified protein, the majority of TorD existed as a monomer, yet dimers and higher order oligomers/aggregates were also observed (Figure 2B, top). The oligomeric state of FPLC-purified monomeric TorD was stable at -80°C for at least a month (Figure 2B, bottom). Considering that the total TorD concentration was significantly higher during initial purification (∼170 µM; Figure 2B, top) than for the combined monomer fractions (∼7 µM; Figure 2B, bottom) or during native gel electrophoresis (∼17 µM; Figure 2A), these data are consistent with a weak and reversible dimerization of TorD (*K*_*D*_ > 50 µM; estimated from the monomer/dimer ratio in Figure 2B).

**Table 1.** Plasmids encoding overproduced proteins.

| Plasmid | Parent | Protein(s) Produced | Comments | Reference |
| --- | --- | --- | --- | --- |
| pTorD-H6 | pET28a | TorD-H6 | C-terminal 6xHis tag | this work |
| pH6-spTorA-mCherry | pET28a | H6-spTorA-mCherry | N-terminal 6xHis tag | this work |
| pH6-TEV-spTorA-GFP(C) | pBAD24 | H6-spTorA-GFP | N-terminal 6xHis tag + TEV protease sequence | this work |
| p-spTorA-GFP-H6C | pBAD24 | spTorA-GFP-H6C | C-terminal 6xHisC tag | (1) |
| p-preSufI(IAC) | pET25b | pre-SufI(IAC) | C17I, C295A, and C-terminal 6xHisC tag | (38) |
| pTatABC | pBAD22 | TatA, TatB, TatC | Used to generate Tat<sup>++</sup> IMVs | (67) |

**Figure 1.**
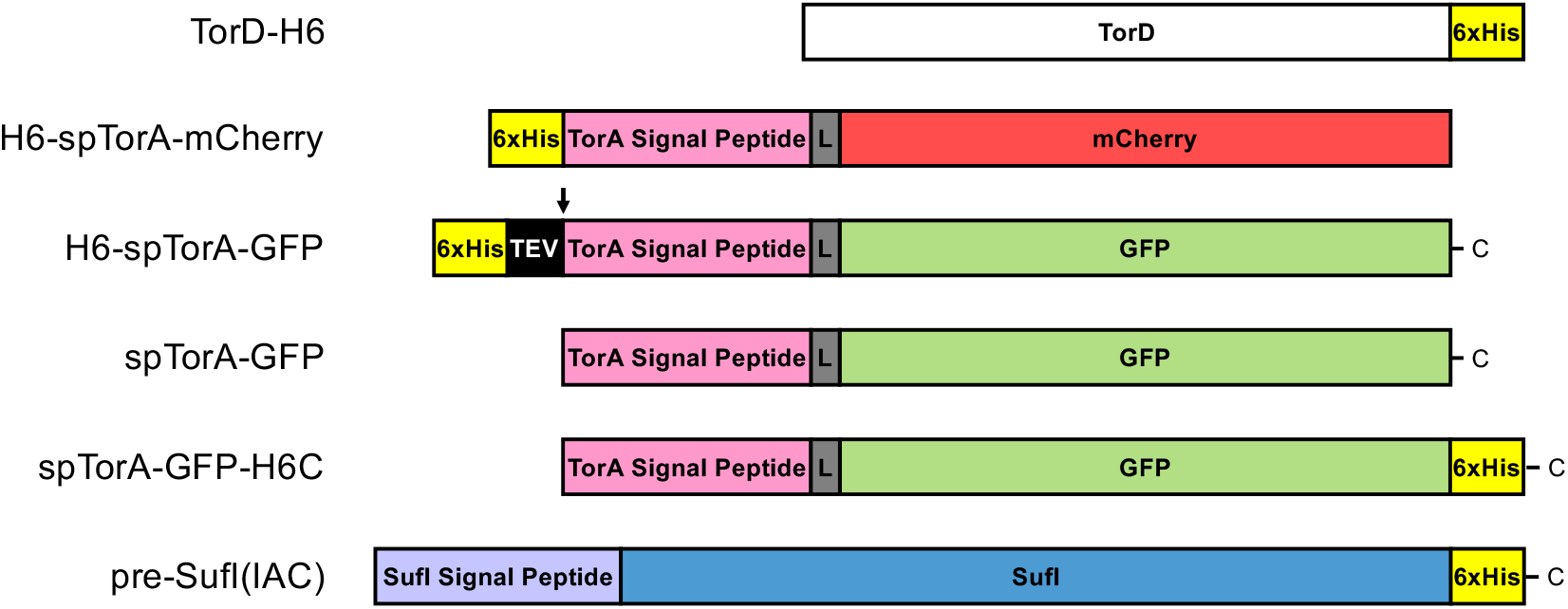
Proteins used in this study. Protein sequences are provided in Figure S1 and plasmid sequences are available from Addgene. A short linker (L) is indicated in *gray*. The TEV protease cleaves within the TEV recognition sequence (ENLYFQG) between Q and G.

**Figure 2.**
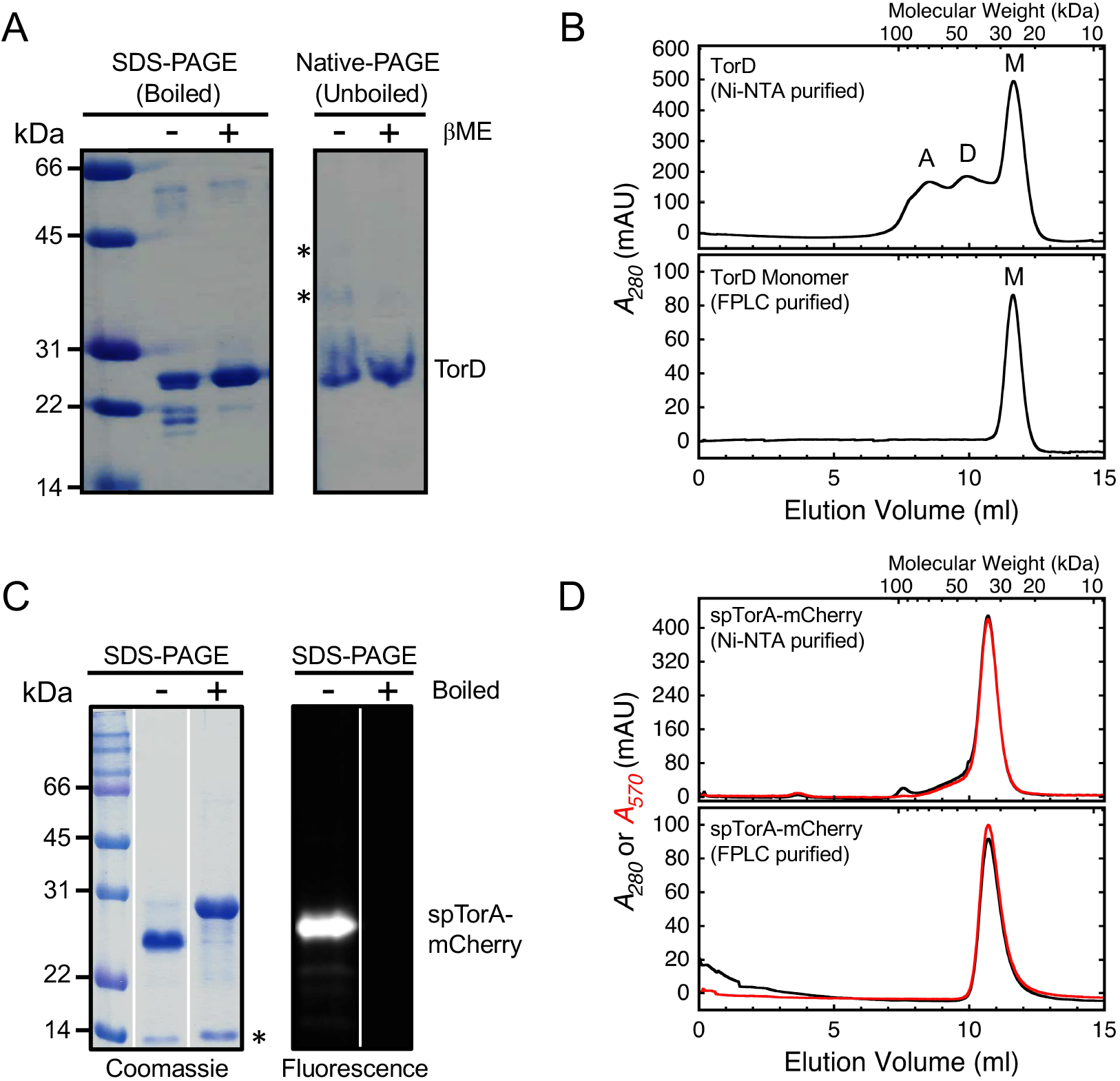
Oligomerization State of TorD and spTorA-mCherry. (A) Oligomerization state of Ni-NTA purified TorD under native gel conditions. TorD is largely monomeric (∼17 µM load), although some higher order oligomers (starred bands) are eliminated by βME (143 mM) and hence are disulfide-linked. (B) Oligomerization state of TorD analyzed by size-exclusion chromatography. Ni-NTA purified TorD contains higher order oligomers at high concentration (*top*, ∼170 µM total load), yet the monomeric form in the FPLC-purified fractions was stable for at least a month at -80°C when preserved in the presence of 50% glycerol and 5 mM DTT at a lower concentration (*bottom*, ∼7.0 µM load). (C) Ni-NTA purified spTorA-mCherry. The fully denatured unfolded form of mCherry (boiled sample) runs slower on SDS-PAGE and is non-fluorescent. The starred (*) band is a heat-dependent cleavage product of mCherry (61). (D) Oligomerization state of spTorA-mCherry analyzed by size-exclusion chromatography. The Ni-NTA purified spTorA-mCherry is largely monomeric (*top*, ∼290 µM total load), and remains stably monomeric upon re-chromatographing (*bottom*, ∼7.1 µM total load). The molecular weight axis on the top of the size-exclusion chromatograms was generated by a standard curve from the peak elution positions of conalbumin (75 kDa), carbonic anhydrase (29 kDa), RNase (13.7 kDa), and aproptinin (6.5 kDa). The ordinates are milli-absorbance units (mAU).

### Monomeric TorD binds to spTorA-mCherry in a 1:1 ratio

We next sought to address whether monomeric TorD is capable of binding to spTorA fused to the fluorescent protein mCherry (spTorA-mCherry; purified and used herein as the 6xHis tagged version H6-spTorA-mCherry). Purified monomeric spTorA-mCherry (Figures 2C & 2D) was mixed with TorD in a 1:1 or 1:2 ratio, incubated at 37°C for 10 minutes, and then analyzed by size-exclusion chromatography. Peaks corresponding to TorD, spTorA-mCherry, and the TorD/spTorA-mCherry complex were readily resolvable, clearly indicating that TorD and spTorA-mCherry formed a 1:1 complex (Figures 3A & 3B). As controls, we examined whether complexes could form between TorD and mCherry or the authentic Tat precursor pre-SufI (purified and used herein as the 6xHis tagged high transport efficiency version pre-SufI(IAC) (38)). Since no such complexes were observed (Figures 3C & 3D), we conclude that TorD specifically recognizes the TorA signal peptide.

**Figure 3.**
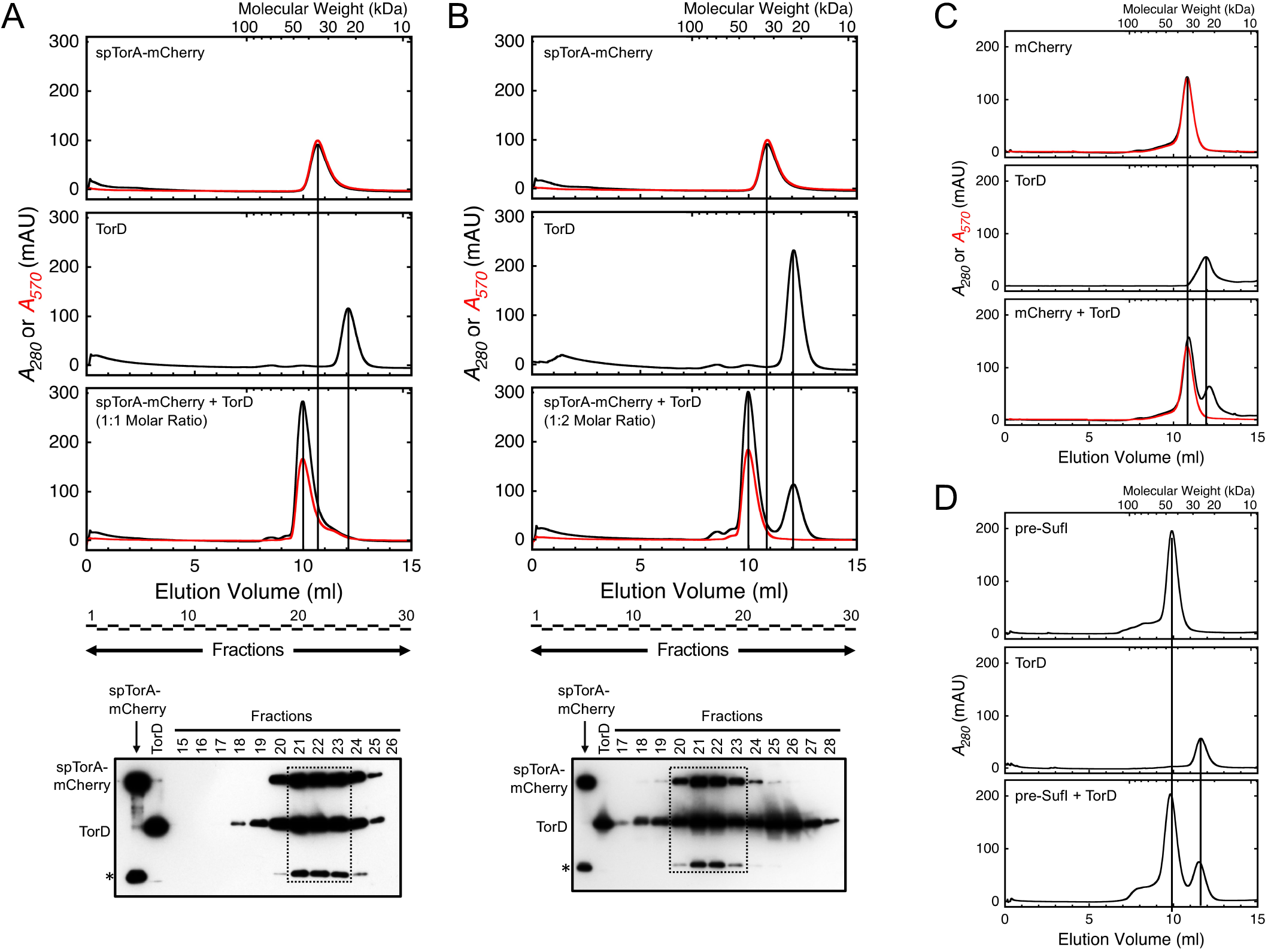
TorD and spTorA-mCherry form a 1:1 complex. (A & B) The TorD/spTorA-mCherry complex. The spTorA-mCherry protein (∼7 µM) was mixed with TorD in a 1:1 (A) or 1:2 (B) ratio and analyzed by size-exclusion chromatography (*bottom chromatograms*). For the 1:1 mixture, the peaks corresponding to the individual spTorA-mCherry (*top chromatogram*) and TorD (*middle chromatogram*) proteins are virtually absent, and the new peak at ∼45 kDa reflects the TorD/spTorA-mCherry complex. For the 1:2 mixture, approximately half of the TorD was recovered uncomplexed with spTorA-mCherry. Integrated signal intensities from Western blots of the elution fractions from the spTorA-mCherry + TorD mixtures confirm a 1:1 stoichiometry of the spTorA-mCherry and TorD proteins in the high molecular weight peak (*dashed boxes*). The lower molecular weight band for purified spTorA-mCherry (*) is a known product of heat-dependent self-cleavage (see text). The spTorA-mCherry and TorD load in the standard lanes was 2 pmol. (C & D) Non-binding controls. TorD does not form a complex with either mCherry, which has no signal peptide, or pre-SufI, which has a non-cognate signal peptide.

To test the stability of the REMP/substrate complex, fractions containing the purified TorD/spTorA-mCherry complex were combined, centrifuged to remove aggregates, and immediately loaded back onto the size-exclusion column. A peak corresponding to monomeric TorD was recovered, indicating partial dissociation of the complex (Figure 4). While a corresponding peak for spTorA-mCherry is expected based on this result, such a peak was not observed. We ascribe the absence of free spTorA-mCherry in this sample to the known tendency of this protein to adhere to surfaces, particularly in the absence of other proteins (such as BSA; data not shown), most likely due to the hydrophobicity of the signal peptide. Partial dissociation of the TorD/spTorA-mCherry complex is expected if the concentration drops significantly and is near the *K*_*D*_. As the total concentration of TorD was ∼2-3-fold higher when the TorD/spTorA-mCherry complex was originally purified (Figure 3A; 7 µM) than when this complex was re-assayed (Figure 4; ∼2.3-3.5 µM), the *K*_*D*_ can be estimated as follows. Since the A_280_/A_570_ ratio is ∼1 for mCherry, the ratio of the two peaks in Figure 4 after subtracting the mCherry absorbance indicates that close to half of the TorD dissociated from spTorA-mCherry, thus indicating that the TorD, spTorA-mCherry, and the TorD/spTorA-mCherry complex concentrations were all approximately 2.3-3.5 µM/2 = 1.2-1.8 µM, or ∼1.5 µM. Assuming a single binding equilibrium where [TorD][spTorA-mCherry]/[TorD•spTorA-mCherry] = *K*_*D*_, the binding affinity is then estimated as (∼1.5 µM)^2^/(∼1.5 µM) = ∼1.5 µM.

**Figure 4.**
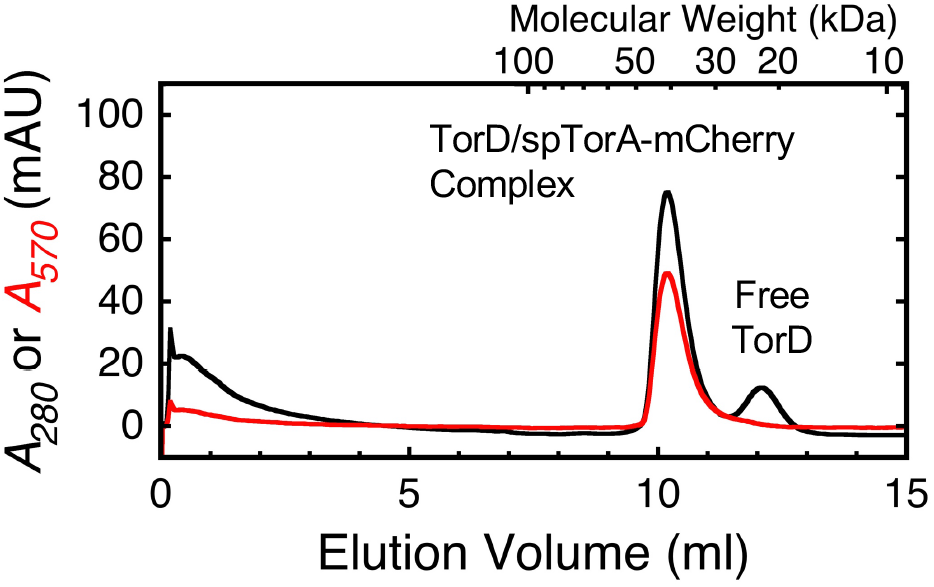
Dissociation of the TorD/spTorA-mCherry Complex. Fractions 19-21 of the 1:1 TorD/spTorA-mCherry complex in Figure 3A were pooled, centrifuged to remove aggregates (4°C, 32,000 *g* for 10 minutes), and immediately re-run on the size-exclusion column. Approximately half of the TorD dissociated from spTorA-mCherry (see text).

### High transport efficiency of spTorA-GFP, a His-tag-free Tat substrate

Cleavage of the signal peptide during purification of Tat substrates is a general problem, typically leading to mixtures of full-length and mature-length proteins (i.e., with and without the signal peptide). Purification of full-length spTorA-mCherry was assured by placing the 6xHis affinity tag at the N-terminus of the protein (Figure 1). However, this location for the 6xHis-tag can potentially interfere with Tat-dependent transport (see later). Moreover, the mCherry protein undergoes an internal heat-dependent self-cleavage (Figures 2C, 3A, & 3B) (59-61), which complicates analysis using SDS-PAGE gels. Therefore, we created H6-spTorA-GFP, which includes a TEV protease site after the N-terminal 6xHis-tag and replaces the mCherry fluorescent protein with GFP (Figure 1). The fluorescent dye Alexa532 was covalently attached to an introduced cysteine at the C-terminus through maleimide chemistry, allowing fluorescence detection on SDS-PAGE after boiling the samples, which destroys the fluorescence of the GFP domain. Removal of the 6xHis-tag by the TEV protease yielded spTorA-GFP(Alexa532) (Figure 5A).

Comparison of *in vitro* Tat transport efficiencies of H6-spTorA-GFP(Alexa532) and spTorA-GFP(Alexa532) revealed that the N-terminal 6xHis+TEV sequence inhibited transport by ∼80% (Figure 5B). The spTorA-GFP(Alexa532) transport efficiency of ∼40% was ∼20% less than the transport efficiency of pre-SufI(Alexa647) (Figure 5B). The transport efficiency of spTorA-GFP(Alexa532) is the best that we have observed for a TorA signal peptide substrate (1,38).

**Figure 5.**
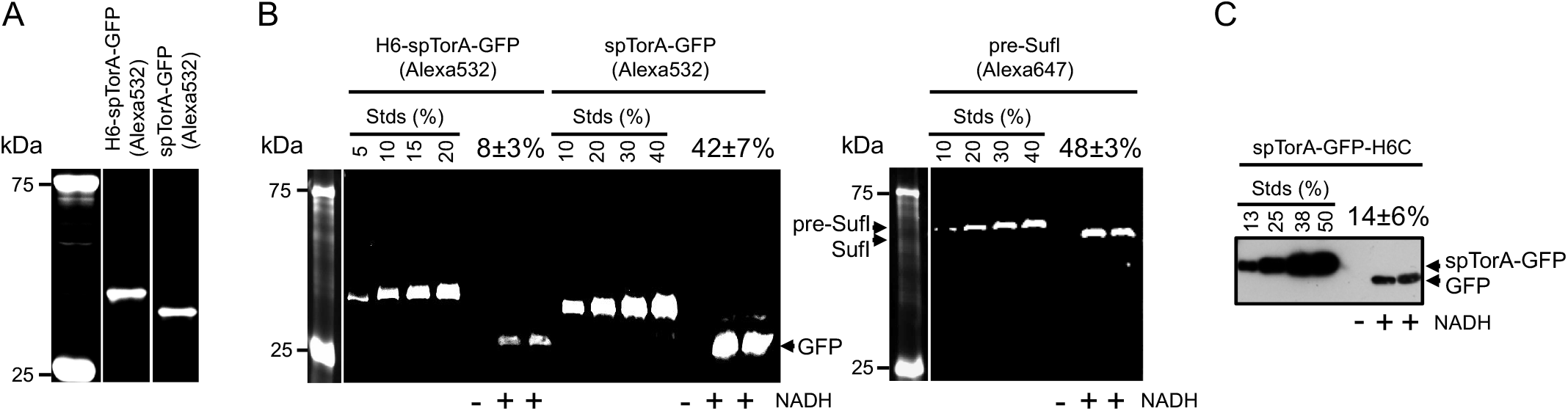
*In vitro* transport efficiencies of Tat substrates. (A) Purified spTorA-GFP(Alexa532) after TEV protease cleavage of H6-spTorA-GFP(Alexa532). (B & C) Transport assays analyzed by in-gel fluorescence (B) or Western blotting (C). Transport assays were conducted with 50 nM precursor proteins and Tat^++^ IMVs (*A*_280_ = 5) for 10 min at 37°C. Transport efficiencies are indicated as the amount of protease-protected (571 µg/mL proteinase K treatment for 20 min) mature-length protein as the percent of total added precursor from at least 4 independent assays. These data suggest that the transport of spTorA-GFP is reduced by ∼5- and 3-fold with a 6xHis-tag at the N- or C-terminus, respectively (but see Figure 6), and that the transport efficiency of spTorA-GFP is ∼80% of that observed for pre-SufI. Transport was not observed in the absence of NADH (control).

### Anti-6xHis Western blots underestimate spTorA-GFP transport efficiency

The Tat transport efficiency of spTorA-GFP-H6C using Western blots with anti-6xHis antibodies was ∼1/3^rd^ of that observed for spTorA-GFP(Alexa532) (Figures 5B and 5C). An ∼5-fold lower transport efficiency was observed for the N-terminally 6xHis-tagged H6-spTorA-GFP(Alexa532) relative to the 6xHis-free spTorA-GFP(Alexa532) protein (Figure 5B). These data suggested that the 6xHis-tag might generally interfere with transport, particularly since the N- and C-termini of GFP are on the same side of the β-barrel structure, and hence the C-terminus of the GFP domain will be near the TatBC receptor complex when it binds to spTorA-GFP. To probe whether the observed transport efficiency differences could be influenced by detection method (chemiluminescence Western blotting vs. in-gel fluorescence), we investigated precursor detection efficiency in the presence and absence of inverted membrane vesicles containing overproduced TatABC (Tat^++^ IMVs). We observed that the in-gel fluorescence detection of spTorA-GFP(Alexa532) was linearly dependent on load and unaffected by the presence or absence of IMVs. In contrast, Western blot detection of H6-spTorA-GFP and spTorA-GFP-H6C was severely underestimated in the presence of IMVs (Figure 6). Poor membrane transfer, detection interference by IMV components, or His-tag cleavage may all contribute to the poor Western detection efficiency (none of these were pursued further). In short, we conclude that poor Western blot detection efficiency of 6xHis-tagged spTorA-GFP proteins by anti-6xHis antibodies in the present of IMVs significantly underestimated the transport efficiencies of these proteins.

**Figure 6.**
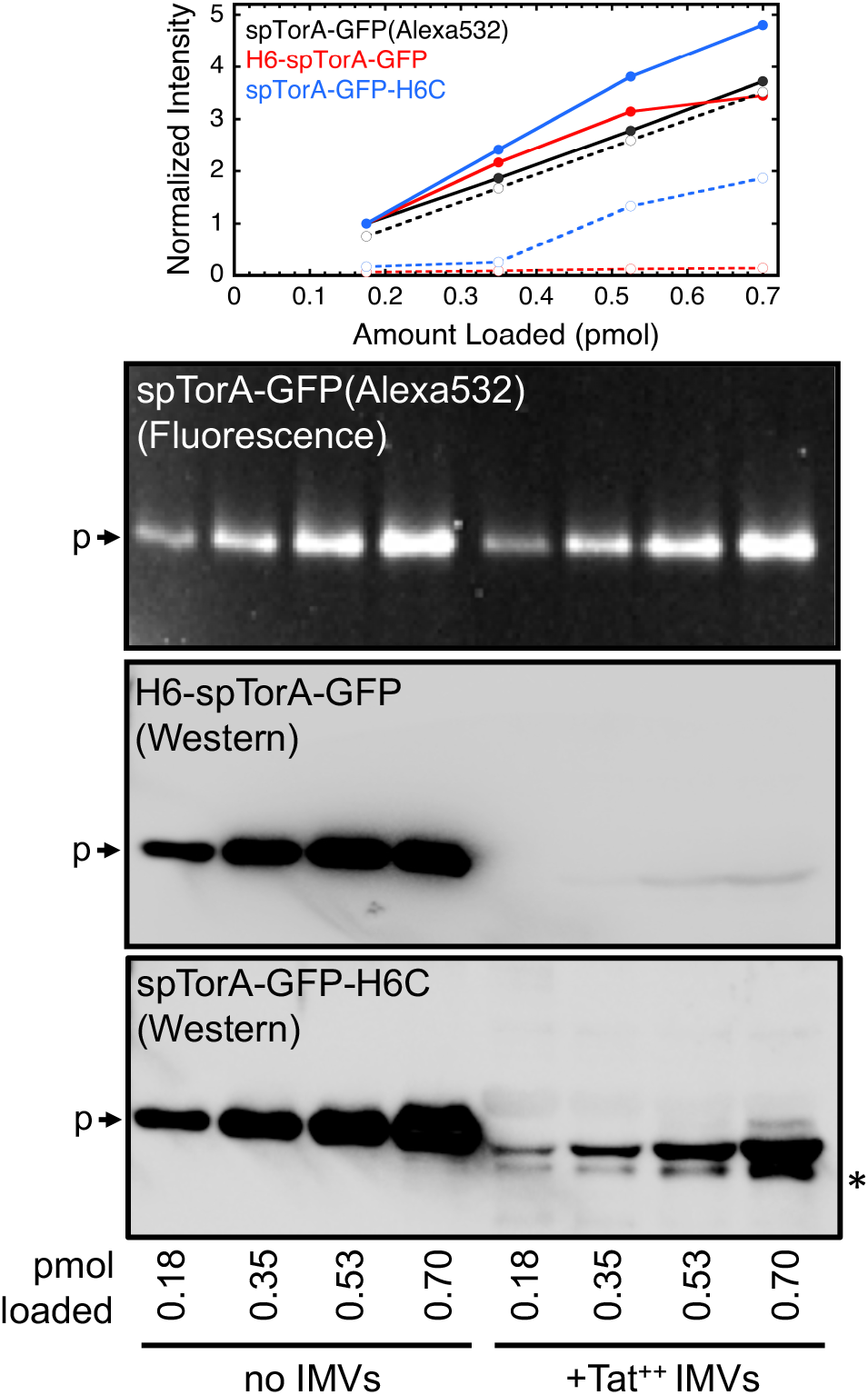
Western blots underestimate Tat precursor concentrations in the presence of Tat^++^ IMVs. Different amounts of the indicated Tat precursors were electrophoresed in the absence or presence of Tat^++^ IMVs (*A*_280_ = 5). In the graph at the top, the intensity dataset for each gel is normalized to the intensity for the 0.18 pmol load in the absence of IMVs: *solid curves*, no IMVs; *dashed curves*, +IMVs. The amount of spTorA-GFP(Alexa532) detected by in-gel fluorescence in the absence or presence of Tat^++^ IMVs is linear with the load, and similar under the two conditions. In contrast, Western blots of H6-spTorA-GFP and spTorA-GFP-H6C using anti-6xHis antibodies substantially underestimate the presence of these proteins when electrophoresed with Tat^++^ IMVs. The starred (*) band indicates a partially cleaved protein product, which is not the mature-length protein.

### TorD inhibits binding of spTorA-GFP to Tat^++^ IMVs

If TorD delivers spTorA-containing substrates to the Tat translocon, the expectation is that TorD would enhance binding of spTorA-GFP to Tat^++^ IMVs. This was not observed. The amount of spTorA-GFP(Alexa532) bound to Tat^++^ IMVs *decreased* with increasing TorD concentrations, with an apparent *K*_*D*_≈ 1.3 µM (Figure 7). This apparent *K*_*D*_ could certainly reflect the affinity of TorD for spTorA-GFP(Alexa532), a reasonable explanation being that TorD bound to the signal peptide prevented the precursor substrate from binding to the TatABC-containing membranes. Alternatively, it may also reflect a spTorA-GFP binding site on the membrane that also binds TorD (competitive binding). Since substrate binding to the membranes was not enhanced by TorD, the binding interactions would need to be mutually exclusive such that substrate binding would be inhibited when binding sites are occupied by TorD. This latter possibility was addressed by examining the binding affinity between TorD and membranes with (Tat^++^) or without (ΔTat) Tat translocons. A similar binding affinity (∼100 nM) was observed for both membranes (Figure 8), suggesting that TorD binds to non-Tat components, most likely the lipid surface. One possibility is that the membrane interaction was mediated by the dye (Alexa532) on TorD. Since the ΔTat and Tat^++^ membranes behaved similarly, these data also argue that any direct binding of TorD to Tat components must be weak, if it occurs. Since the TorD affinity for membranes is substantially stronger than the TorD inhibition of substrate binding to the Tat^++^ IMVs (Figure 7), the most reasonable interpretation is that the *K*_*D*_ ≈ 1.3 µM reflects the binding affinity of TorD for the TorA signal peptide after the membrane binding sites are saturated with TorD, and that the TorD/spTorA-GFP complex does not bind to IMVs. Note that this apparent *K*_*D*_ (≈ 1.3 µM) is similar to the *K*_*D*_ (≈ 1.5 µM) for the TorD/spTorA-mCherry complex estimated from the FPLC chromatogram (Figure 4; see earlier discussion).

**Figure 7.**
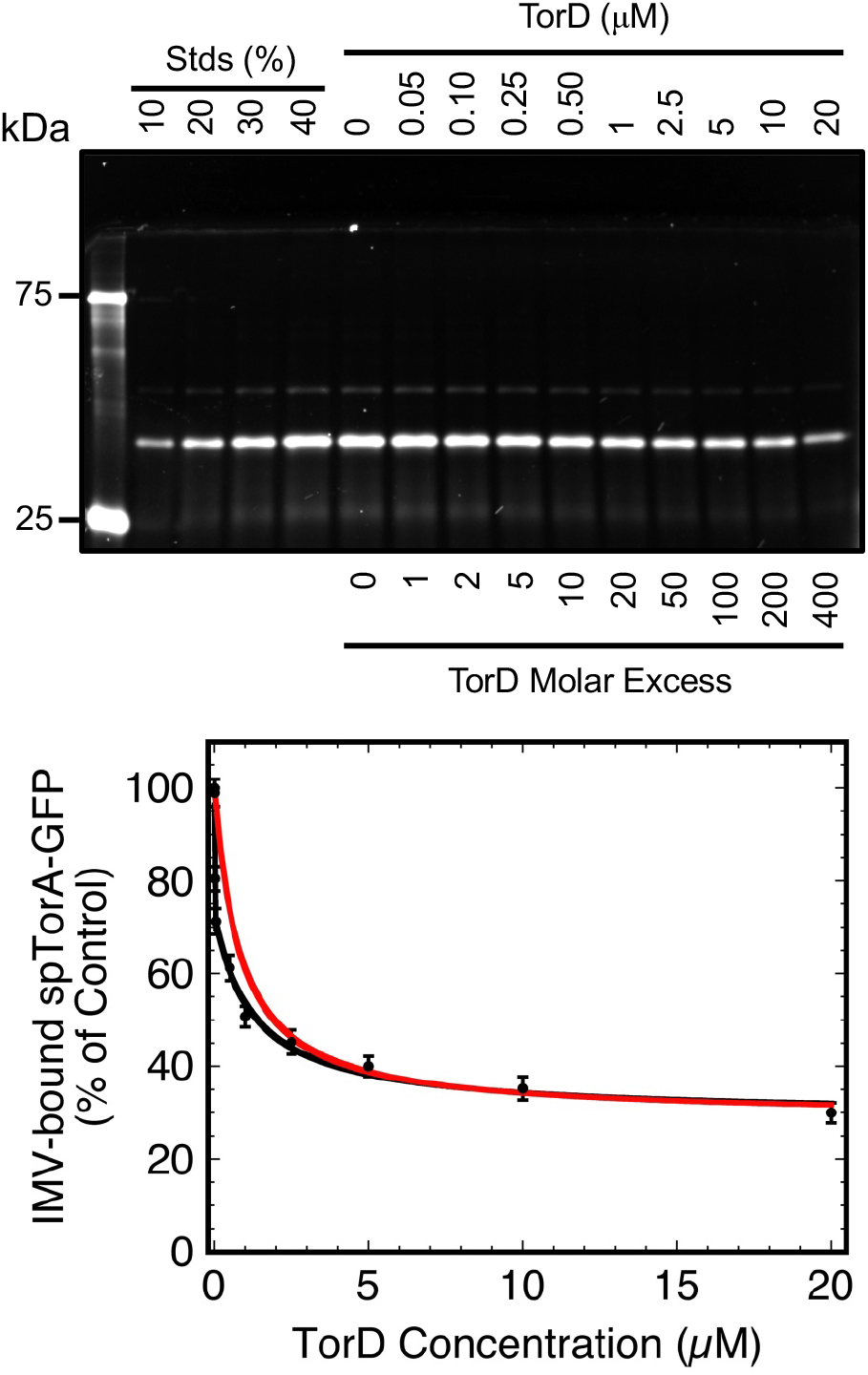
TorD reduces the binding of spTorA-GFP to Tat^++^ IMVs. The spTorA-GFP(Alexa532) protein (50 nM) was pre-incubated with various concentrations of TorD (10 min, 37°C), incubated with Tat^++^ IMVs (*A*_280_= 5; 10 min at 37°C), and centrifuged to remove unbound protein (32,000 *g*, 4°C for 10 min). The pellets were analyzed by SDS-PAGE, and the amount of IMV-bound spTorA-GFP(Alexa532) was quantified by in-gel fluorescence imaging using the standard lanes for calibration (% of 50 nM). Data are normalized to the 0 µM TorD sample (100%; *N* = 3). Assuming that the TorD interaction with spTorA-GFP prevents the precursor from binding to the IMVs, three binding regimes are apparent: first phase (∼30%), linear fit; second phase (∼40%), single site Langmuir binding isotherm (*K*_*D*_ = 1.3±0.3 µM) (46); third phase (∼30%), no apparent binding to TorD. The first two phases (independent fits in *black*) are also simultaneously fit to a single site binding model that accounts for the precursor concentration, but not the IMV binding sites (*red curve*) (31).

**Figure 8.**
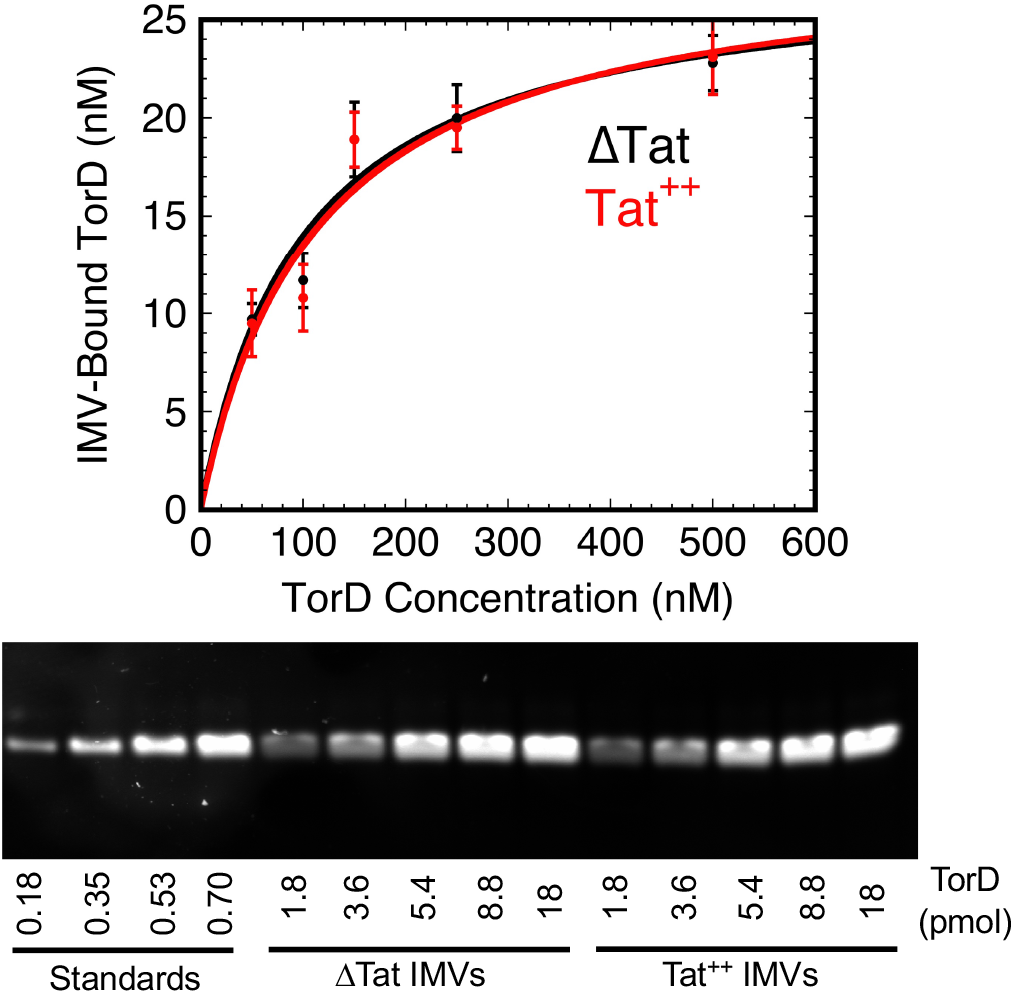
TorD has similar affinities for ΔTat and Tat^++^ IMVs. Different concentrations of TorD(Alexa532) were incubated with ΔTat and Tat^++^ IMVs (*A*_280_ = 5; 10 min at 37°C). IMV pellets were recovered and analyzed for the amount of bound TorD using the approach described for Figure 7. Data were fit using a single site Langmuir binding isotherm model (46), yielding *K*_*D*_ values of 100±33 nM and 112±43 nM as well as [TorD]_bound(max)_ values of 28±3 nM and 29±4 nM for ΔTat and Tat^++^ IMVs, respectively.

### TorD minimally inhibits transport of spTorA-GFP

Tat-dependent transport of spTorA-GFP was performed under the same conditions as the membrane binding assay, except that NADH was added to generate the pmf needed for transport (Figure 9). While the transport efficiency at 20 µM TorD was reduced to 30% of the TorD-free control, similar to the reduction in membrane binding by TorD (compare Figures 9B and 7B), the former decrease was linear and the latter was logarithmic. These data therefore indicate that the effect of TorD on binding and transport occur due to distinctly different phenomena. Importantly, while membrane binding of spTorA-GFP was ∼50% inhibited by ∼1 µM TorD, this TorD concentration affected transport efficiency by < 5%.

**Figure 9.**
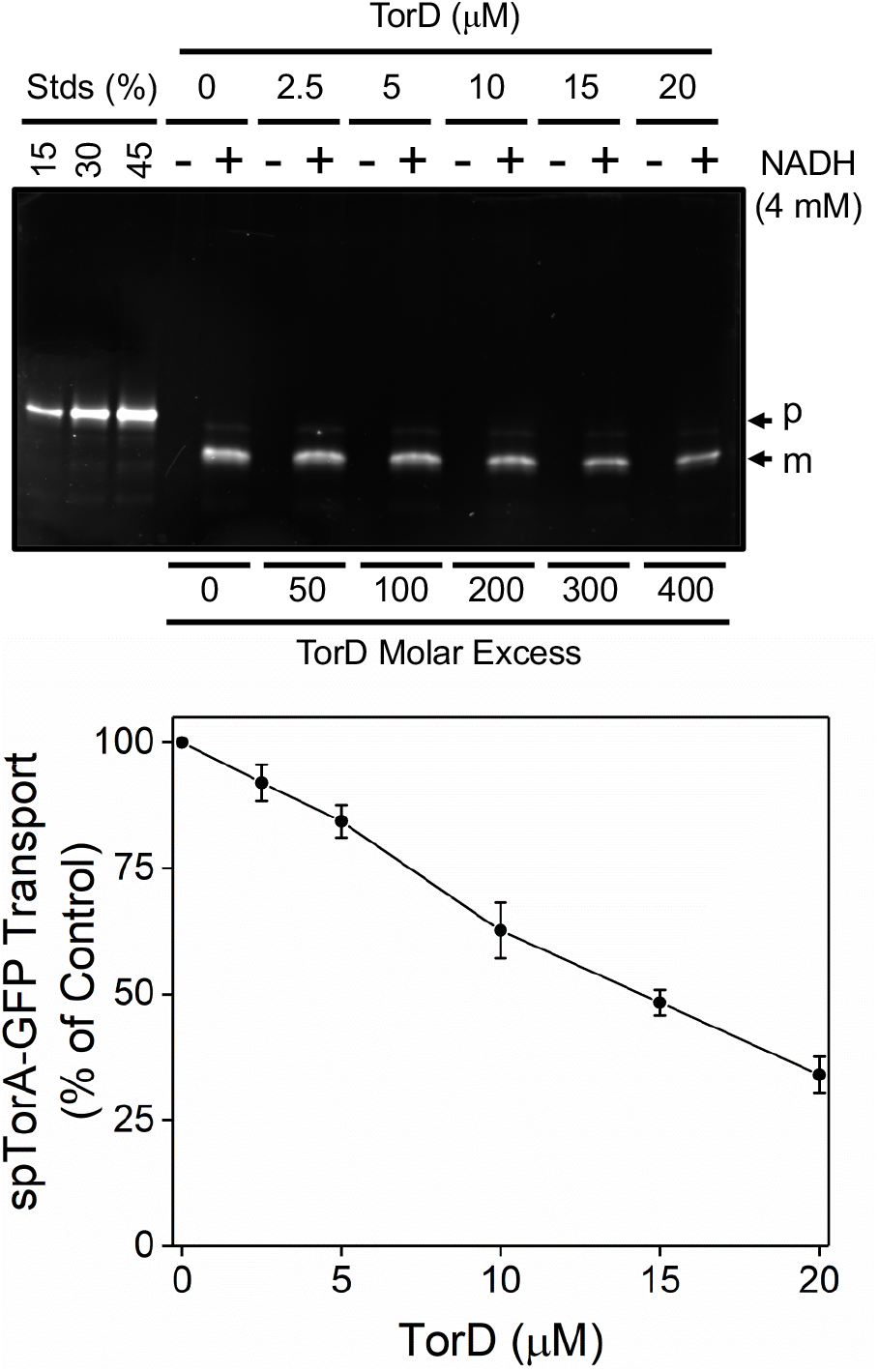
spTorA-GFP transport is inhibited by TorD. Tat-dependent transport of spTorA-GFP(Alexa532) (50 nM) was measured in the presence of increasing concentrations of TorD (conditions of Figure 5). Markers were not used for this experiment since all lanes were used for the assay. The standard lanes provide a suitable reference (*N* = 3).

## DISCUSSION

This study tested the hypothesis that REMPs could enhance Tat transport efficiency by migrating with their cognate substrates to the Tat translocon and then hand-off the signal peptide from the REMP binding site to the Tat receptor complex. Our major findings are as follows: 1) TorD is largely monomeric at low micromolar concentrations; 2) the TorD/spTorA interaction has a *K*_*D*_ of ∼1-2 µM; 3) TorD binds to the cytoplasmic membrane, but this does not mediate any enhanced substrate interaction with this membrane; 4) TorD does not bind to any Tat structural proteins; and 5) TorD does not enhance Tat-dependent transport, but rather minimally inhibits transport. These findings are consistent with a model in which TorD and the spTorA-containing substrates used here are in rapid dynamic equilibrium, and only the REMP-free form of the substrate binds to the Tat receptor complex to initiate the transport process.

*Sewanella massilia* TorD is dimeric under conditions similar to Figure 2A, and its X-ray structure revealed an extreme domain swapped dimer (54,55). However, monomeric *E. coli* TorD binds the TorA signal peptide (45), raising questions about the active form of TorD *in vivo*. A domain swapped dimer is not expected to readily interconvert between dimer and monomer forms during normal physiological processes. We found here that the *E. coli* TorD oligomerizes, but that this is concentration dependent with a dimerization *K*_*D*_ in excess of 50 µM, the dimers dissociate upon dilution, and purified monomeric TorD is stable. We also found that monomeric TorD has a micromolar affinity for spTorA, and the interconversion between bound and unbound state is sufficiently fast that it does not substantially interfere with Tat-dependent transport. Thus, for an *in vivo* TorD concentration in the range of tens of micromolar or less, we expect that TorD serves its function as a monomer.

The three-phase titration curve of the IMV-substrate binding interaction with increasing amounts of TorD (Figure 7) indicates heterogeneity. The most likely explanation is distinct signal peptide conformations that do not readily interconvert and that differentially interact with TorD. In this experiment, the spTorA-GFP substrate was pre-incubated with TorD before adding IMVs, so the precursor protein certainly had the opportunity to bind to TorD unhindered by membranes. Nonetheless, ∼30% of the precursor protein bound to the IMVs at high concentrations of TorD and did not track the TorD/spTorA affinity, and thus this population of precursor protein is considered non-interactive with TorD. The remaining ∼70% of the spTorA-GFP precursor protein was inhibited from binding to IMVs by TorD in two distinct phases (*black fit* in Figure 7). In the first phase (∼30%), the TorD binding interaction appears to be approximately linear due to a very strong binding interaction. In the second phase, the data are well fit by a single site Langmuir binding isotherm with a *K*_*D*_ ≈ 1.3 µM. These two phases together are decently fit by a binding model that accounts for the precursor concentration (*red fit* in Figure 7), yielding an approximate *K*_*D*_ of ∼0.8 µM; however, this model does not account for precursor binding to the IMV membrane, and therefore must be considered only an approximation. Together with the estimate for the TorD/spTorA-mCherry affinity from the size-exclusion analysis (∼1.5 µM; Figure 4), all three approaches consistently reveal a TorD/spTorA interaction strength of ∼1 µM. This is consistent with the high end values from previous results, which range from 0.06-4 µM (16,45,46,48). The previously determined extreme high affinity value is consistent with the first binding phase in Figure 7.

The apo (cofactor-free) pre-TorA precursor has a mature domain site that interacts with TorD with sufficient affinity such that a TorD/apo-TorA complex (no spTorA or molybdopterin cofactor present) can be isolated (47). This suggests a bivalent interaction between TorD and apo-pre-TorA that is substantially stronger than the ∼1 µM *K*_*D*_ for the TorD interaction with the signal peptide alone. A stable, high affinity TorD/apo-pre-TorA interaction protects the signal peptide from degradation (11,18), and also prevents targeting to/recognition by the Tat translocon. TorD binds the mature molybdopterin cofactor and assists in delivering it to apo-pre-TorA to generate the properly assembled holo-enzyme (10,12,13). The trigger for release of TorD from pre-TorA is most likely proper insertion of the molybdopterin cofactor, which results in a weakening of the TorA mature domain interaction with TorD (13). According to this picture, the interaction of the fully assembled holo-enzyme pre-TorA likely interacts with TorD much the same as spTorA-GFP does, that is, largely via the signal peptide alone since the TorA mature domain has a weakened interaction with TorD. Thus, we expect that the effects of TorD on the membrane binding and transport efficiency of spTorA-GFP reported here similarly apply to fully-assembled pre-TorA.

While TorD does bind to IMVs, we have no evidence for any TorD interaction with the Tat translocon in the presence or absence of the spTorA-GFP substrate. Therefore, this study argues against the hypothesis that REMPs target substrates to the Tat translocon. While REMP interactions with their cognate mature domains could potentially significantly modulate the strength of signal-peptide interactions as well as interactions with the Tat translocon, we favor the simpler model described earlier in which proper cofactor insertion leads to distinctly weaker REMP interactions with their holo-enzyme substrates. We therefore conclude that REMPS do not promote Tat-dependent transport at the level of the translocon, though by protecting signal peptides during substrate folding and assembly, they can ensure a greater transport yield of synthesized proteins.

## EXPERIMENTAL PROCEDURES

### Bacterial strains, growth conditions, and plasmids

The *E. coli* strains MC4100ΔTatABCDE, JM109, and BL21(λDE3) were described earlier (62-64). Overexpression cultures were grown in Luria-Bertani (LB) medium at 37°C supplemented with appropriate antibiotics (65). All plasmids overproducing the proteins described in Figure 1 that were constructed by us were submitted to Addgene, and the construction of new plasmids is described in the history of the linked SnapGene files. All coding sequences were verified by DNA sequencing. The construction of the three novel plasmids reported here is briefly outlined below, and the encoded amino acid sequences are indicated in Figure S1.

#### pTorD-H6

The TorD-6xHis coding sequence was amplified from pQE80TorADhis (47) and inserted into pET28a using NcoI and HindIII restriction sites. The asparagine mutation at position 46 was converted back to the wildtype serine by inverse PCR.

#### pH6-spTorA-mCherry

The spTorA-mCherry-6xHisC coding sequence was amplified from p-spTorA-mCherry-H6C (66) and inserted into pET28a using NcoI and HindIII restriction sites. Limited digestion was used as there is an NcoI restriction site within mCherry. This internal NcoI site was then removed by the QuikChange protocol (Agilent Technologies). The 6xHis tag was switched to the N-terminus using PCR amplification and the fragment was inserted back into pET28a with NcoI and a filled-in and blunted HindIII site.

#### pH6-TEV-spTorA-GFP(C)

The internal NcoI site within the GFP coding sequence of p-spTorA-GFP-H6C (1) was eliminated by PCR amplification, and the amplified fragment was inserted back into p-spTorA-GFP-H6C using NcoI and MscI restriction sites. Then, a 6xHis tag and TEV sequence were added to the N-terminus of spTorA-GFP and the 6xHis tag was removed from the C-terminus using PCR amplification, and the amplified fragment was inserted back into p-spTorA-GFP-H6C using NcoI and PstI restriction sites.

### Protein production and purification

The TorD, spTorA-mCherry, and pre-SufI proteins were overproduced in BL21(λDE3) in 2 l conical flasks and purified under native conditions by Ni-NTA chromatography. LB cultures (500 ml) were shaken at 200 rpm, 37°C until the *A*_600_reached ∼3. The pH of the cultures was raised to ∼9.0 by the addition of 25 ml 0.5 M CAPS (pH 9.0), and protein production was induced with 1 mM IPTG for 2.5 h. The TorD and pre-SufI cultures were induced at 37°C, and the spTorA-mCherry cultures were induced at 25°C. Cultures were chilled on an ice bath and centrifuged at 5,000 g for 12 min at 4°C. Pellets were rapidly resuspended on ice in 50 ml Buffer A (100 mM Tris, 25 mM CAPS, pH 9.0) containing 1 M NaCl, 0.2% Triton X-100 with protease inhibitors (10 mM PMSF, 100 µg/ml trypsin inhibitor, 20 µg/ml leupeptin, and 100 µg/ml pepstatin), DNase (20 µg/ml), and RNase (10 µg/ml). Cells were passed through a French pressure cell once at 16,000 psi. The cell lysate was cleared of cellular debris by centrifugation at 50,000 *g* for 30 min at 4°C, and then stirred with 2 ml Ni-NTA Superflow resin (Qiagen) that had been pre-equilibrated with Buffer A for 10 min on ice. The resin was loaded onto a 10 x 1 cm column, and sequentially washed with: (1) 100 ml of Buffer B (10 mM Tris-HCl, 1 M NaCl, pH 8.0) with 0.1% Triton X-100; (2) 20 ml of Buffer B; (3) 20 ml of Buffer C (10 mM Tris-HCl, 100 mM NaCl, pH 8.0); and (4) 10 ml of Buffer C containing 50% glycerol, pH 8.0. Proteins were eluted with Buffer C with 500 mM imidazole, 50% glycerol, pH 8.0 in 1 ml fractions, and stored at -80°C. Typical yield was 12–20 mg of total protein per liter of culture.

The spTorA-GFP-H6C and H6-spTorA-GFP proteins were overproduced in MC4100ΔTatABCDE after induction with 1% arabinose. The spTorA-GFP-H6C protein was purified under denaturing conditions and refolded by dilution/dialysis from 9 M urea as described earlier (1). The H6-spTorA-GFP protein was purified under native conditions using Ni-NTA chromatography. LB cultures (1000 ml) were shaken at 200 rpm, 37°C until the *A*_600_ reached ∼1.5-2. The pH of the cultures was raised was raised to ∼9.0 by the addition of 50 ml 0.5 M CAPS (pH 9.0), and protein production was induced (1% arabinose) at 25°C for ∼12 h. The culture was chilled on an ice bath and centrifuged at 5,000 *g* for 12 min at 4°C. Pellets were rapidly resuspended on ice in 50 ml Buffer A containing 1X CelLytic B (Cat. #C8740, Sigma, MO, USA), and the suspension was incubated for 10 min on ice with occasional mixing. The cell lysate was cleared of cellular debris by centrifugation at 50,000 *g* for 30 min at 4°C. The supernatant was mixed with 3 ml Ni-NTA Superflow resin that had been pre-equilibrated with Buffer A containing 1X CelLytic B for 10 min on ice. The resin was loaded onto a 10 x 1 cm column, and the H6-spTorA-GFP protein was washed, eluted and stored as described in the previous paragraph. Typical yield was 3–5 mg of protein per liter of culture.

### Labeling of purified proteins with fluorescent dyes

Ni-NTA purified proteins were labeled on cysteines with fluorescent dyes for easier visualization within polyacrylamide gels. Proteins (∼25 µM in 100 µl) were incubated with 1 mM *tris*[2-carboxyethylphosphine] hydrochloride (TCEP) for 10 min, and then treated with Alexa532 or Alexa647 maleimide (Invitrogen) for 15 min at room temperature in the dark. The dye excess required for quantitative labeling was determined by titrating the dye to protein ratio to determine the point of labeling saturation. A 20-fold excess was required for TorD(Alexa532) and pre-SufI(Alexa647), whereas a 50-fold excess was used to produce H6-spTorA-GFP(Alexa532). Reactions were quenched with 10 mM βME and purified by Ni-NTA chromatography essentially as explained earlier (38). Labelled proteins were diluted ∼50-fold with Buffer C to reduce the concentration of imidazole to 10 mM or lower, and then mixed with 0.4 ml of Ni-NTA Superflow resin pre-equilibrated with Buffer C. The resin was loaded onto a 3×0.5 cm column and washed with 10 mL of Buffer C, and then with 5 mL of Buffer C containing 50% glycerol. The labelled precursor was eluted (0.2 ml fractions) with 100 mM NaCl, 500 mM imidazole, 50% glycerol, pH 8.0, and stored at -80°C. Typically, the labeling efficiency was > 90%, as determined from SDS-PAGE after Coomassie Blue staining since both labeled and unlabeled proteins were resolved.

### Purification and analysis by size-exclusion chromatography

Size-exclusion chromatography was performed using an AKTAdesign FPLC system (Amersham Pharmacia Biotech). The peak fractions of spTorA-mCherry and TorD eluted from Ni-NTA resin were ∼200-400 µM. In each case, 400 µl of protein solution was diluted with 100 µl FPLC buffer (5 mM MgCl_2_, 150 mM NaCl, 25 mM MOPS, 25 mM MES, 25% glycerol, pH 8.0) supplemented with 5 mM DTT, and then incubated on ice for 10 min. After centrifugation to remove any insoluble aggregates (32,000 g, 4°C for 10 min), the supernatant was loaded onto a Superdex 75 10/300 GL size-exclusion column and run with FPLC buffer at a flow rate of 0.5 ml/ min at 4°C. Collected fractions (0.5 ml) were supplemented with an equal volume of 10 mM DTT, 50% glycerol in FPLC buffer and stored at -80°C.

To examine the affinity between spTorA-mCherry and TorD, the two proteins were mixed in a 1:1 or 1:2 ratio and incubated at 37°C for 10 min in FPLC buffer supplemented with 5 mM DTT. Oligomerization was analyzed by size-exclusion chromatography as described for their purification in the previous paragraph. Aliquots (5 µL) from elution fractions were analyzed by Western blotting. The TorD binding interactions with mCherry and pre-SufI were analyzed identically.

### Western blotting

PVDF membranes were used for Western blotting. All steps (membrane blocking, primary antibody treatment, secondary antibody treatment and washing steps to remove loosely bound antibodies to membrane) were performed at room temperature in Western buffer (1X PBS (137 mM NaCl, 2.7 mM KCl, 10 mM Na_2_HPO_4_, 2 mM KH_2_PO_4_, pH 7.4) supplemented with 1% BSA, 0.1% Triton X-100, and 0.1% Tween 20). PVDF membranes were blocked (1 h) with Western buffer prior to adding primary antibodies. To detect 6xHis-tagged proteins, blocked membranes were incubated (1 h) first with mouse anti-6xHis polyclonal antibodies (1:5000; Santa Cruz Biotechnology, Inc.) and then with rabbit anti-mouse antibodies conjugated with HRP (1:10,000; Invitrogen, Inc). Each antibody incubation was followed by two 5 min wash steps.

### Formation and purification of spTorA-GFP(Alexa532)

H6-spTorA-GFP(Alexa532) (200 µl, 10-15 µM) was dialyzed in a 1 ml dialysis cup (10 kDa cutoff) against TEV cleavage buffer (50 mM Tris-HCl, 0.5 mM EDTA, 1 mM DTT, pH 8.0) at 4°C for 2 h to substantially exchange the buffers. TEV protease (100 units) was added, and the dialysis was continued for ∼12 h. The contents from the dialysis cup were quantitatively recovered by puncturing the membrane and centrifuging into a fresh microfuge tube. While the TEV-cleaved N-terminal tag (6xHis+TEV recognition peptide) was largely removed through dialysis, complete removal of the tag and any uncleaved protein was ensured by incubating the dialysate with 100 µl of Ni-NTA equilibrated with TEV cleavage buffer on ice for 30 minutes with periodic mixing. The contents were centrifuged and the supernatant was preserved by adding glycerol and 250 mM DTT in 50X TEV cleavage buffer to yield a final concentration of 5 mM DTT and 48% glycerol.

### Isolation of inverted membrane vesicles (IMVs)

IMVs were isolated essentially as described previously (1), with a few modifications. The spheroplast formation buffer was altered by increasing the concentration of EDTA to 2 mM and the lysozyme concentration to 0.8 mg/ml. After incubation (20 min on ice), the suspension was diluted 4-fold to reduce the EDTA concentration. The spheroplasted cells were passed through a French Press at 12,000 psi, as compared to the originally described 6,000 psi. The MC4100ΔTatABCDE strain required a much higher pressure for optimal formation of IMVs, as compared to JM109 cells. In addition, the 2.2 M sucrose cushion was replaced with a 3-step (0.5, 1.5, and 2.3 M) sucrose gradient, which enabled enrichment of a highly active inner membrane fraction (31). The Tat^++^ and ΔTat IMVs were obtained from MC4100ΔTatABCDE cells that did or did not overproduce TatABC, respectively, as previously described (66).

### Analytical methods

Protein concentrations were determined by the densitometry of bands on SDS-PAGE gels stained with Coomassie Blue R-250 using carbonic anhydrase as a standard and a ChemiDoc MP imaging system (Bio-Rad Laboratories). Fluorescent proteins were detected by direct in-gel fluorescence imaging using the same ChemiDoc imaging system. Alexa532 and Alexa647 concentrations were determined using ε_531_ = 81,000 cm^−1^ M^−1^ and ε_650_ = 270,000 cm^−1^ M^−1^, respectively. Western blot bands were visualized by chemiluminescence using the Clarity Max Western blotting kit (Bio-Rad Laboratories) and the ChemiDoc imaging system. IMV concentrations were determined as the *A*_280_ in 2% SDS (1). All error bars are standard deviations.

### Membrane binding and *in vitro* translocation assays

For membrane binding and *in vitro* translocation assays, we used standard 35 µL reactions, as previously described (1,38). For the membrane binding studies, IMVs (*A*_280_ = 5) and precursor (50 nM) were combined (35 µl reaction volume) in high-BSA translocation buffer (HB-TB), which contains a 10-fold higher concentration of BSA (570 µg/ml) than translocation buffer (TB; 5 mM, MgCl_2_, 50 mM, KCl, 200 mM sucrose, 57 µg/ml BSA, 25 mM MOPS, 25 mM MES, pH 8.0). The high BSA concentration minimizes non-specific binding. Protein LoBind microfuge tubes (1.5 ml, Eppendorf) were used to further minimize non-specific binding to the walls of the reaction vessel. For translocation assays, the pH was 8.0, as this higher pH promotes more efficient transport (38).

## DATA AVAILABILITY

Plasmids used and sequences are available from Addgene. All remaining data are contained within this manuscript.

## ACKNOWLEDGEMENTS

We thank T. Palmer for MC4100(DE3), MC4100ΔTatABCDE and pQE80-TorADhis and T. Yahr for providing pTatABC. This research was supported by the National Institutes of Health (GM116995 to SMM). The content is solely the responsibility of the authors and does not necessarily represent the official views of the National Institutes of Health.

## ABBREVIATIONS

Tat: twin-arginine translocation
REMP: redox enzyme maturation protein
Pmf: proton motive force
IMVs: inverted membrane vesicles
βME: β-mercaptoethanol
TECP: *tris*[2-carboxyethylphosphine] hydrochloride
DTT: dithiothreitol
BSA: bovine serum albumin

---

**Figure S1.**
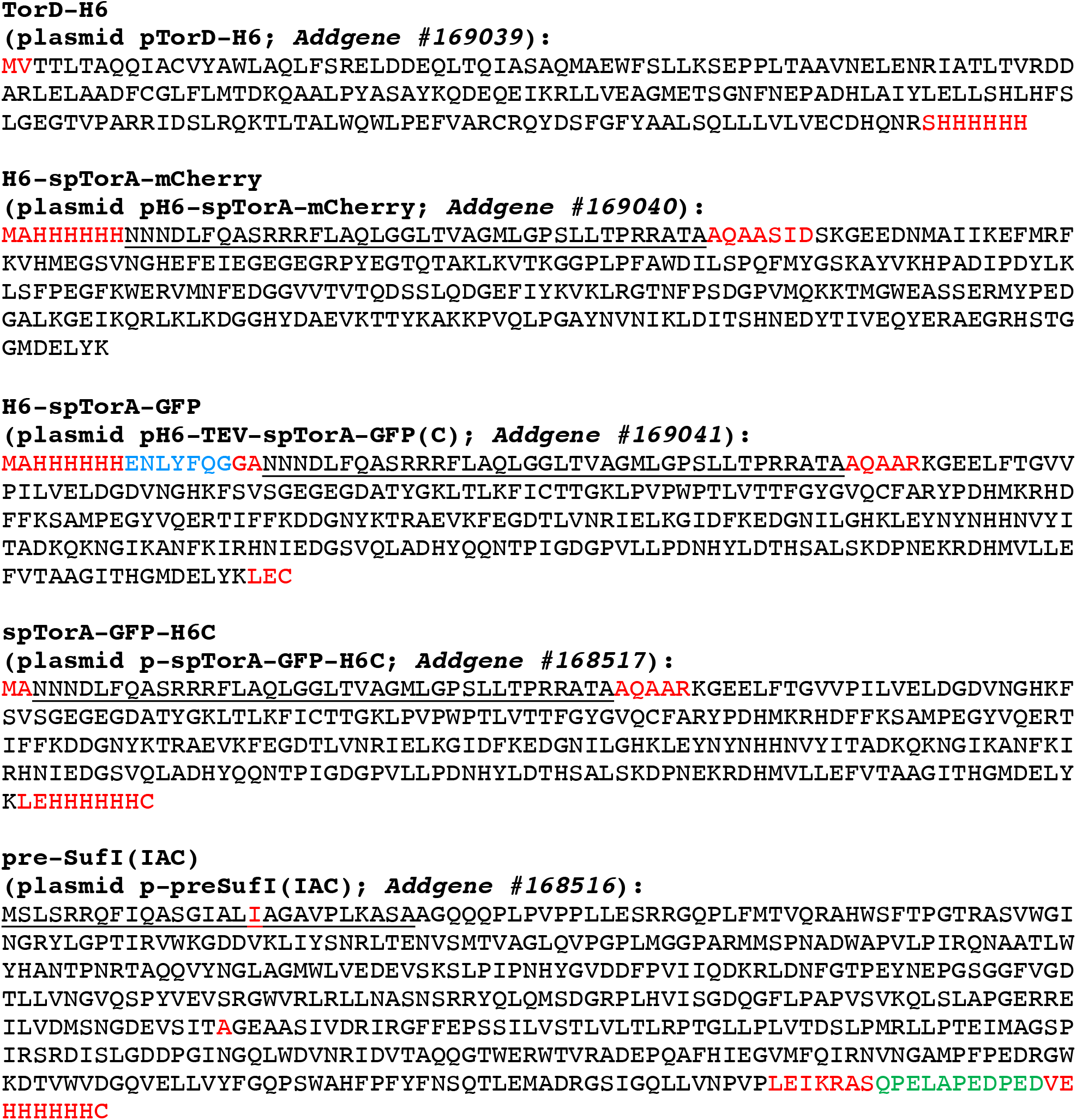
Protein sequences for purified proteins used in this study. The signal peptides of TorA and SufI are underlined, the TEV sequence is identified in *blue*, the QSV tag is identified in *green*, and other additions/linkers/mutations are indicated in *red*.

